# Edge Detection of Cryptic Lamellipodia Assisted by Deep Learning

**DOI:** 10.1101/181263

**Authors:** Chuangqi Wang, Shawn Kang, Eunice Kim, Xitong Zhang, Hee June Choi, Aaron Choi, Kwonmoo Lee

## Abstract

Cell protrusion plays important roles in cell migration by pushing plasma membrane forward. Cryptic lamellipodia induce the protrusion of submarginal cells in collective cell migration where cells are attached and move together. Although computational image analysis of cell protrusion has been done extensively, the study on protrusion activities of cryptic lamellipodia is limited due to difficulties in image segmentation. This study seeks to aid in the computational analysis of submarginal cell protrusion in collective cell migration by using deep learning to detect the cryptic lamellipodial edges from fluorescence time-lapse movies. Due to the noisy features within overlapping cells, the conventional image analysis algorithms such as Canny edge detector and intensity thresholding are limited. In this paper we combined Canny edge detector, Deep Neural Networks (DNNs), and local intensity thresholding. We were able to detect cryptic lamellipodial edges of submarginal cells with high accuracy from the fluorescence time-lapse movies of PtK1 cells using both simple convolutional neural networks and VGG-16 based neural networks. We used relatively small effort to prepare the training set to train the DNNs to detect the cryptical lamellipodial edges in fluorescence time-lapse movies. This work demonstrates that deep learning can be combined with the conventional image analysis algorithms to facilitate the computational analysis of highly complex time-lapse movies of collective cell migration.

## Introduction

Computational analyses of fluorescence images of migrating cells have been used as a major method for quantitative understanding of cell migration^1,2^. Cell migration is a process in which actin filaments are rapidly assembled and disassembled to make the leading edge of the cell to protrude and move forward in respond to its environment^3^. Results of cell migration serve as a great biological importance and are essential in physiological and pathophysiological processes such as wound healing, embryonic development, immune responses, and cancer metastasis.

In both physiological and pathophysiological conditions many cell types migrate together in a group^4^. This collective cell migration plays very important roles in tissue remodeling and morphogenesis since it keeps the tissue intact and allows the communication between cells^4-8^. Collective cell migration shares many mechanistic characteristics with single cell migration, which requires mechanochemical cycles of leading edge protrusion, front adhesion formation, rear adhesion disruption, and cell body contraction^9-11^. However, understanding single cell migration is not sufficient enough to understand collective migration since cells are connected and they communicate with each other. One important characteristic of epithelial sheet migration is the coordination of movement between neighboring cells^12,13^. These cells affect neighboring cells mechanochemically through cadherin-mediated cell-cell junctions and continuously rearrange tissue structures^14-20^. Recent studies showed that cell-cell junctions during collective migration is mechanosensitive^14,15,20^, further suggesting that cells harness mechanical forces to communicate with each other, and the force generation from this cell protrusion process can affect mechanosensitive cell-cell junctions^14^. This suggests that the interaction between cell protrusion and cell-cell junctions are crucial in collective cell migration. To better understand these phenomena, it is essential to develop a quantitative live cell imaging approach to allow us to characterize the coordination of cellular activities among neighboring cells.

Fluorescence imaging has been used as a major method to help capture cell migration patterns^1^. Images taken from the time-lapse recording can then be used to further study important cellular activities in cell migration. Computational image analysis is the way of interpreting information extracted by a computer from a given image^21^, and edge detection is one of the most fundamentals algorithms in image processing. Moreover, it is usually the first step for computational image analysis. Although conventional segmentation method is efficient in identifying outside borders of cells, it is difficult to successfully identify cell edges when they overlap with each other during collective cell migration^22^. Any other noisy image features surrounding the edges could also decrease the accuracy and reliability of the image analysis. While edges of cells are often distinguishable with the human eye, mistakes are often made. As the need to detect edges of cells and patterns in cell migration continues to increase in complexity such as analyzing clusters of cells or whole tissues, a computational method, rather than a manual method, is required^23^.

Deep learning can help by using computers to accurately and efficiently self-determine the edges of cells in a given image, thus helping in analyzing cell migration patterns ^24,25^. Recent progresses in deep learning have shown that computers can outperform humans in analyzing complex, large and/or high dimensional data sets^26-28^. While deep learning has also been applied for image segmentation and edge detection^24,25^, most of the applications are completely independent of the traditional image algorithms such as Canny detector^29^ and intensity thresholding^30^. In this study, we focused on aiding the conventional algorithms using deep learning. We demonstrate that deep learning can be effectively integrated into the conventional image algorithms to enhance the performance of the detection of edges of cryptic lamellipodia.

## Methods and Materials

Fluorescence images of PtK1 cells (kidney epithelial cells from a rat kangaroo) stained with CellMask Orange (Invitrogen) were taken using a spinning disk confocal microscope (Fig. 1A). During their wound healing process, the PtK1 cells tend to attach together and move as a group. As a result, images taken of PtK1 cells can contain cell boundaries of multiple cells, which may make image analysis difficult. Also, the cryptic lamellipodial protrusion is underneath the adjacent cells, meaning that we need to segment the cryptic lamellipodial protrusion from the background of other cells (Fig. 1A). This background cell images are highly variable in terms of images features and signal intensity. Therefore, it is very difficult to develop a segmentation algorithm relying on conventional image analysis algorithm. To tackle this complexity, we established a segmentation pipeline combining deep neural networks (DNN) with other image analysis algorithms as described in Fig. 2.

**Figure 1.**
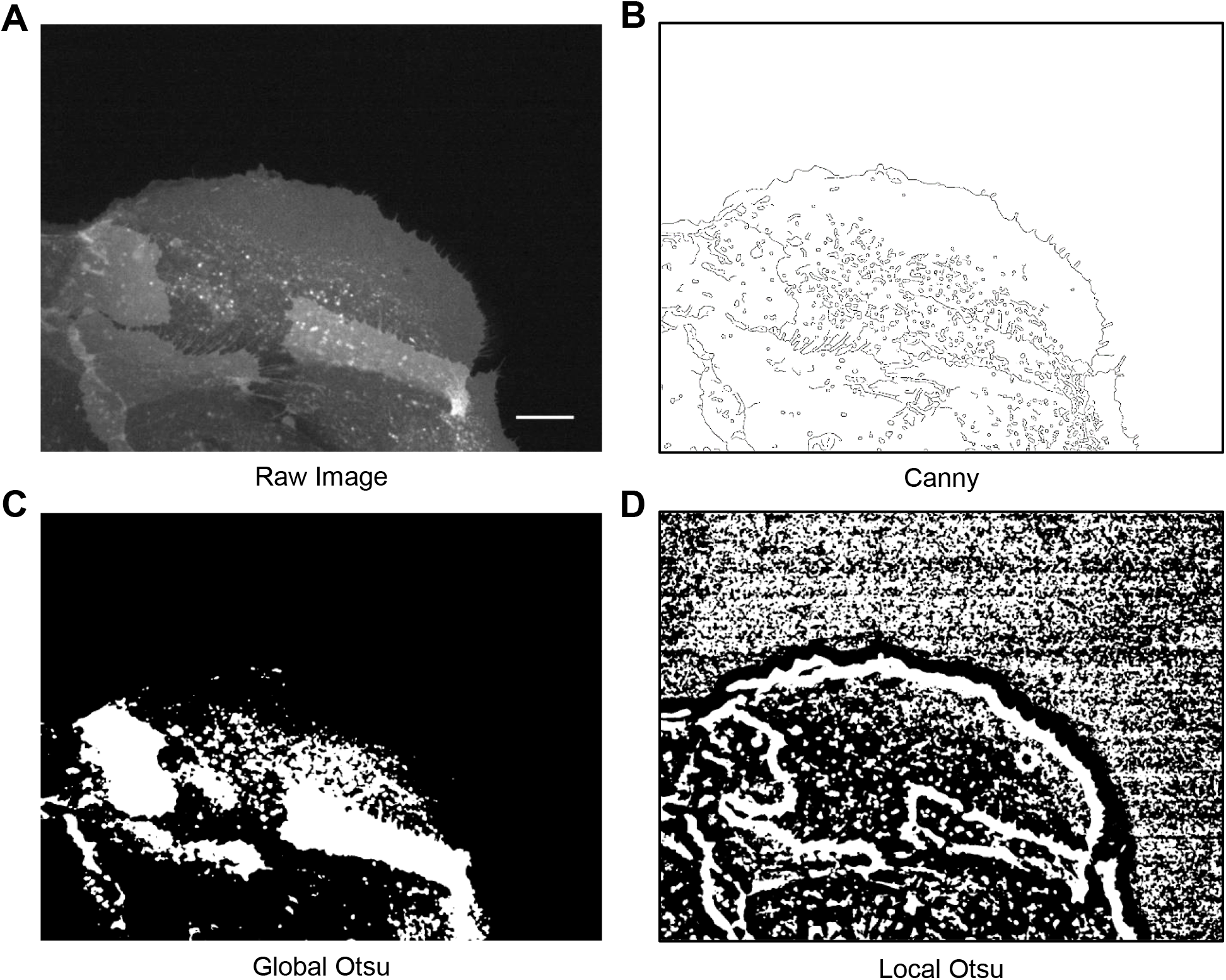
Results of conventional image analysis algorithms for cryptic lamellipodial edges (**A**) A spinning disk confocal image of PtK1 cells stained with a plasma membrane marker, CellMaskOrange^TM^. The image contains multiple PtK1 cells, one whose edge is overlapped by the other cells. Bar: 15mm (**B**) Canny edge image from the image of (A). (**C-D**) The thresholded image of the image of (A) using global (C) and local (D) Otsu algorithms.

**Figure 2.**
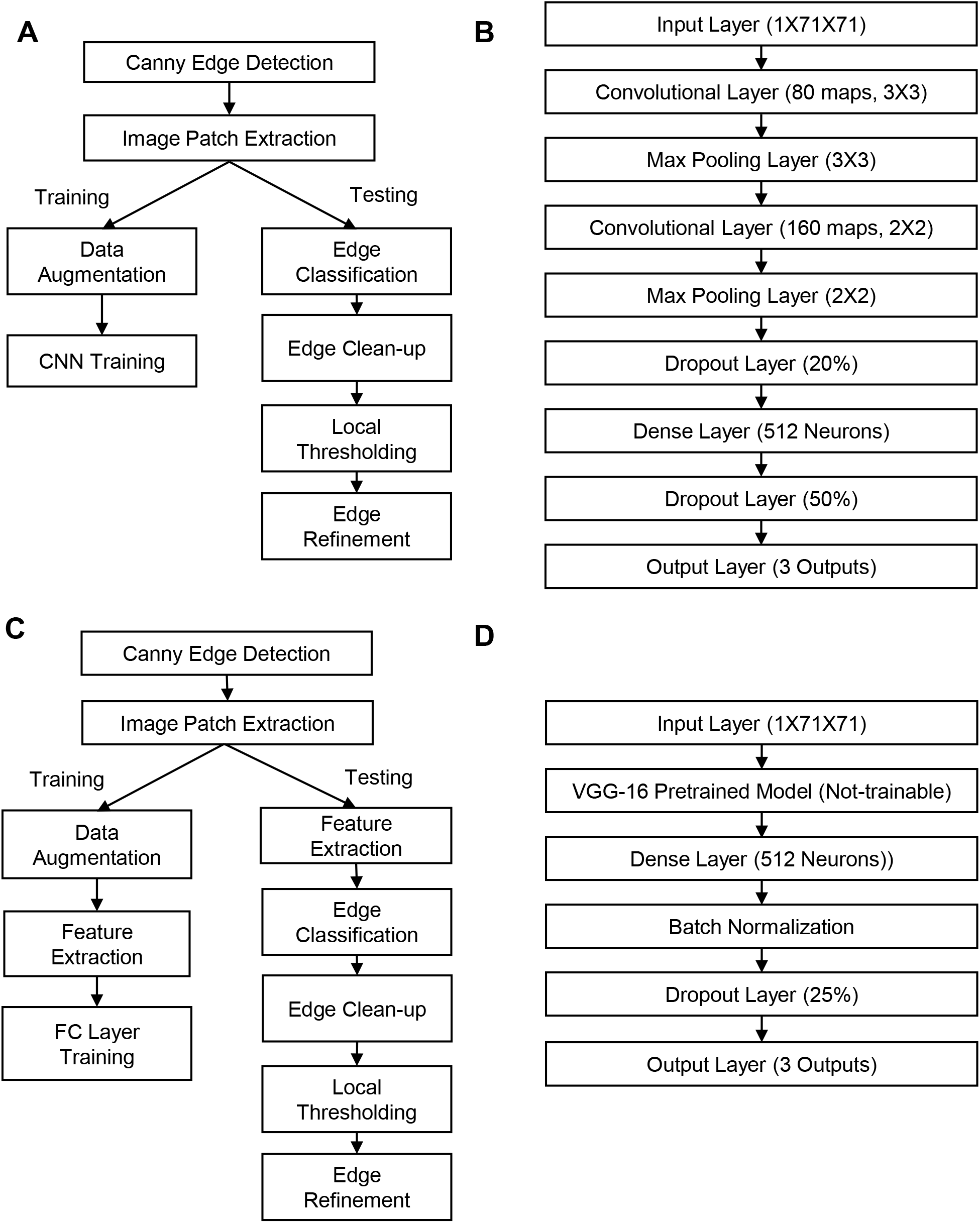
A schematic of the proposed pipeline. (**A**) Computational pipeline for the inner edge detection combining Canny edge detector, CNN, and local thresholding. (**B**) Deep learning architecture in (A). **(C)**The pipeline using VGG-16 pretrained model (VGG-16- DNN). (**D**) Deep learning architecture in (C)

Our pipeline starts with first selecting possible cell-edge candidates through Canny edge detection^29^. By focusing on these edge candidates, we can substantially reduce the computational cost. The Canny algorithm begins by using Gaussian smoothing to remove noise in the image. The edge filters, which calculate the image gradient are used to determine the possible location of edges. Then, non-maximum suppression is used to make the edges thinner^29^. The resulting “Canny image” contained enhanced edges of the original cell image that served as possible edge candidates (Figure 1B).

These edge candidates included the outer and inner boundaries of the cell, which served as real edges of the cell, as well as the other numerous noisy edges inside the cell, which were originated from other cellular features. These false positive edge candidates needed to be removed, and outer and inner edges needed to be distinguished to create a cell image with distinct outer and inner edges. To achieve this, we trained Deep Neural Networks (DNNs) based on convolutional neural networks^31^. The input data of DNNs are small image patches (71X71 pixels) whose center has the edge candidate point from Canny edge detector. The output of the DNNs are the classification results of outer/inner/noisy edges. The DNN accepts the raw intensity input from the images and classifies the edges.

In this paper, we used two kinds of DNN structures. First, we used simple Convolutional Neural Networks (CNNs). These CNNs are composed of three types of layers. These layers are the convolutional layers, pooling layers, and fully connected layers. In convolutional layers, filters are applied to each input layer. As the filter is moved across the layer, outputs known as feature maps are produced. In between convolutional layers, there are usually pooling layers. These layers help to reduce overfitting by reducing the size of feature representations. To prevent overfitting, the dropout layer was also used^32^. Finally, at the end of the network, fully connected layers are used to make predictions and create any final feature combinations. As seen in the CNN architecture in Figure 2B, the structure starts with an input layer. The size of input image patches was specified 71x71 pixels. The number of feature maps and the filter size for the next layer, the convolutional layer, was listed in the Fig. 2B. Next, a dropout layer set to randomly take out 20% of the neurons present in the layer was included to prevent overfitting^32^. The following layer, the max pooling layer, was configured to have a pool size of 3x3 and also serves as a way to control overfitting. The flattening layer, which enables the output to be processed by fully connected layer, was then included. The structure ends with a fully connected layer, which has 512 neurons and the output layer. Training was done based on a categorical cross-entropy loss function. Using the result of the loss function, weights were updated through back propagation^33^ to increase the accuracy percentage by training the neural networks to reduce the amount of errors made.

Usually, DNNs have many model parameters, causing the model to overfit the training data. To prevent this, it is required to prepare a large number of training set. In order to mitigate this issue, we also tested the second DNN architecture in this study, VGG-16 model which was pre-trained on the ImageNet Database^34^. This model was used to extract the features from our microscopy images and these features were used for training fully connected neural network (Fig. 2C-D).

To prepare the training set, we chose the images for the training set every 30 frames in the time lapse movie. In this study, the frame rate of the movie was 5 sec/frame, and the movie contained 200 frames. We chose the frames 1, 31, 61, 91 for the training sets. The canny edge images of those frames were created, and the outer, inner, and other edges were manually isolated. Since we relied on the canny detector for this purpose, the human effort for the training set is relatively low in comparison to other edge detection training where humans need to draw the accurate edges pixel by pixel. Three separate images were then created from the markings, and from each image (Fig 3A). Then small image patches (Figure 3B) were extracted around the edge points and marked as either a noisy edge (0), an inner edge (1), or an outer edge (2) depending on whether the center of the image was a part of these boundaries or not. The corresponding coordinates of these patches were matched to the original image to mark where in the original image points of the outer boundary, inner boundary, and of neither boundary existed. The original image was then cropped to create the training set. We used four images out of 100 frame movies to extract total 20,000 images patches for the training set. These data set was randomly divided into 80% training and 20% validation sets. In order to prevent overfitting, we performed the data augmentation before the training. The training data were augmented to 10 times within the ranges of 30 degree rotation and 20% zoom using Keras package.

Once the DNNs had completed training from the training set, a separate cell image was applied for testing. A final image representing the outer and inner edges of the newly applied cell image was then produced based on the trained DNNs. Due to the errors in initial Canny edge detection and the DNN classification, the outcomes from the neural network tend to contain the disconnected edges. To address this issue, we applied the local thresholding to achieve the continuous edges. First, we removed the small isolated edges whose pixel size is less than 5. Subsequently, from each edge point, we extracted the local image patches (61X61) and we calculated the intensity threshold values by DNN. In order to reduce the effects of noise, we first applied Gaussian filter to the original images. Then, we calculated the median edge intensity of the filtered images, which was used to segment the images. After that, we also perform the edge refinement using image opening operation to remove finger-like edges caused by the bright spots.

## Results

CellMaskOrange^™^ stains plasma membrane and provide a convenient way to monitor cellular leading edge dynamics by a spinning disk confocal microscope. High contrast images of leading edges of marginal PtK1 cells in wound healing responses can be obtained (Fig. 1A). However, the leading edges of submarginal cells, which is driven by cryptic lamellipodia, are overlapping with other neighboring cells, which generate the non-uniform background. Also, bright fluorescence puncta make it even more challenging to identify the inner edges of submarginal cells (Fig. 1A). We applied conventional Canny edge detection algorithm to this image. Canny edge detection^29^ can not only identify outer and inner cell edges, but it also generates numerous noisy edges from the inside of the cell. Another common method of image segmentation is intensity thresholding. When Otsu algorithm^30^ is applied to the cell images, it fails to segment the submarginal cells correctly due to the highly non-uniform background (Fig. 1C). To handle this, we also applied local Otsu method^35^ which calculates the local threshold values to segment the images. The result in Fig. 1D suggests that this local thresholding can be a feasible method to identify the inner edges since it correctly identifies cell edges. However, the method can be sensitive to the parameter of local window size. It is necessary to manually fine-tune the parameter in each frame of the time-lapse movie. Therefore, there are inconsistent segmentation results in different frames.

To address this, we combined deep learning technique with Canny edge detection and local thresholding (Fig. 2A). Deep learning can identify true edges from Canny detection results, which will be fed into local thresholding to produce accurate edge detection. The DNNs were trained to classify outer, inner, and noisy edges (Fig. 3). The training curve of the CNN (Fig. 4A) showed that the training accuracy and the validation accuracy increased and the training loss and validation loss decreased (Figure 4A). As demonstrated in the training curves, our CNN model was able to distinguish outer, inner, and other edges, which created outer/inner edge as expected (Fig. 4B). Using the pretrained VGG-16 marginally increased the validation accuracy (Fig. 4C), and the testing results showed that the edge images contained substantially less small noisy edges than the simple CNN model (Fig. 4D).

**Figure 3.**
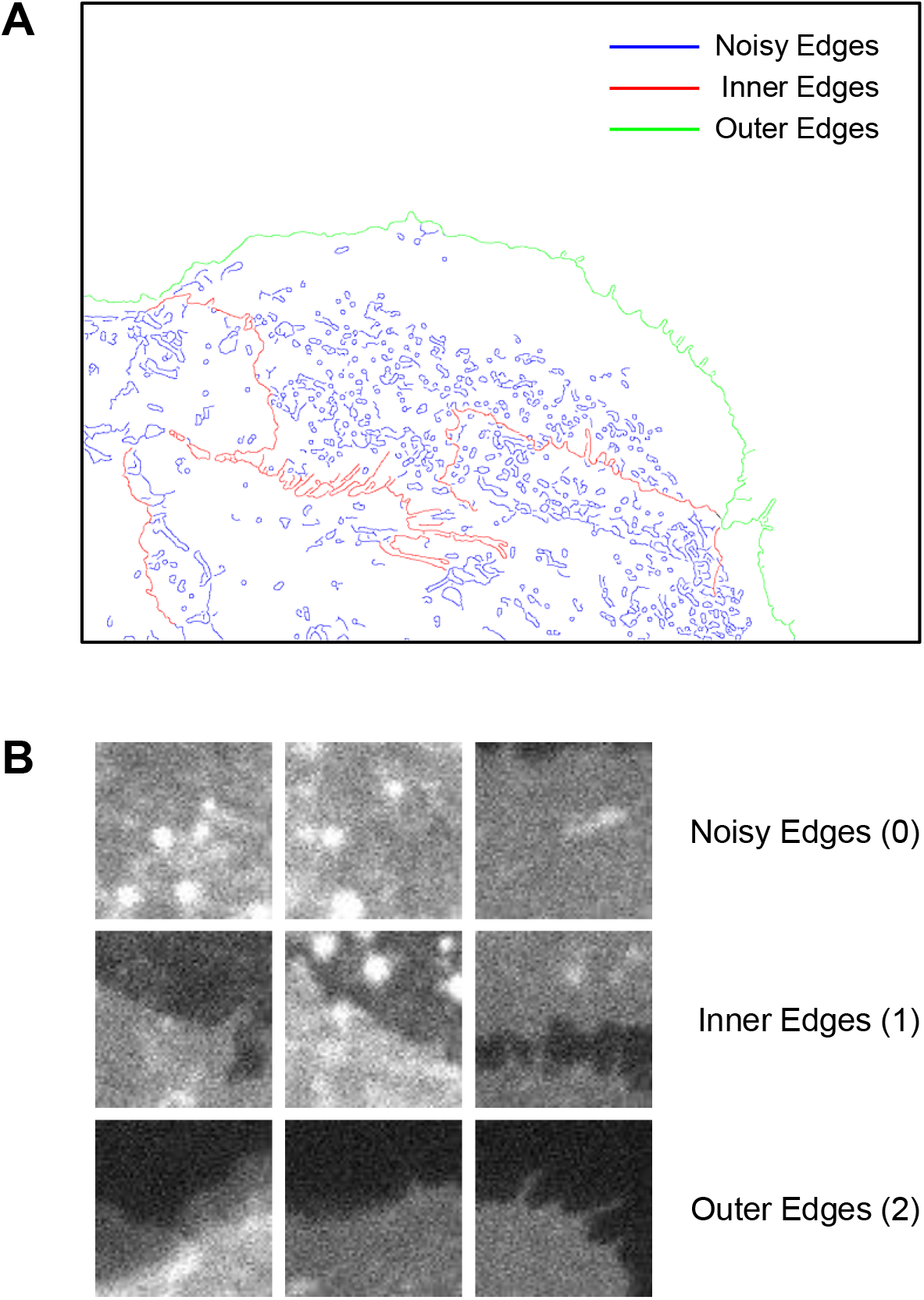
Preparation of the training set. (**A**) The Canny Image was manually divided into three separate images of the outer edges (red), the inner edges (green), and noisy edges (blue). (**B**) The examples of the training set consisted of images patches from the raw cell image that were labeled according to their central points at (36,36).

**Figure 4.**
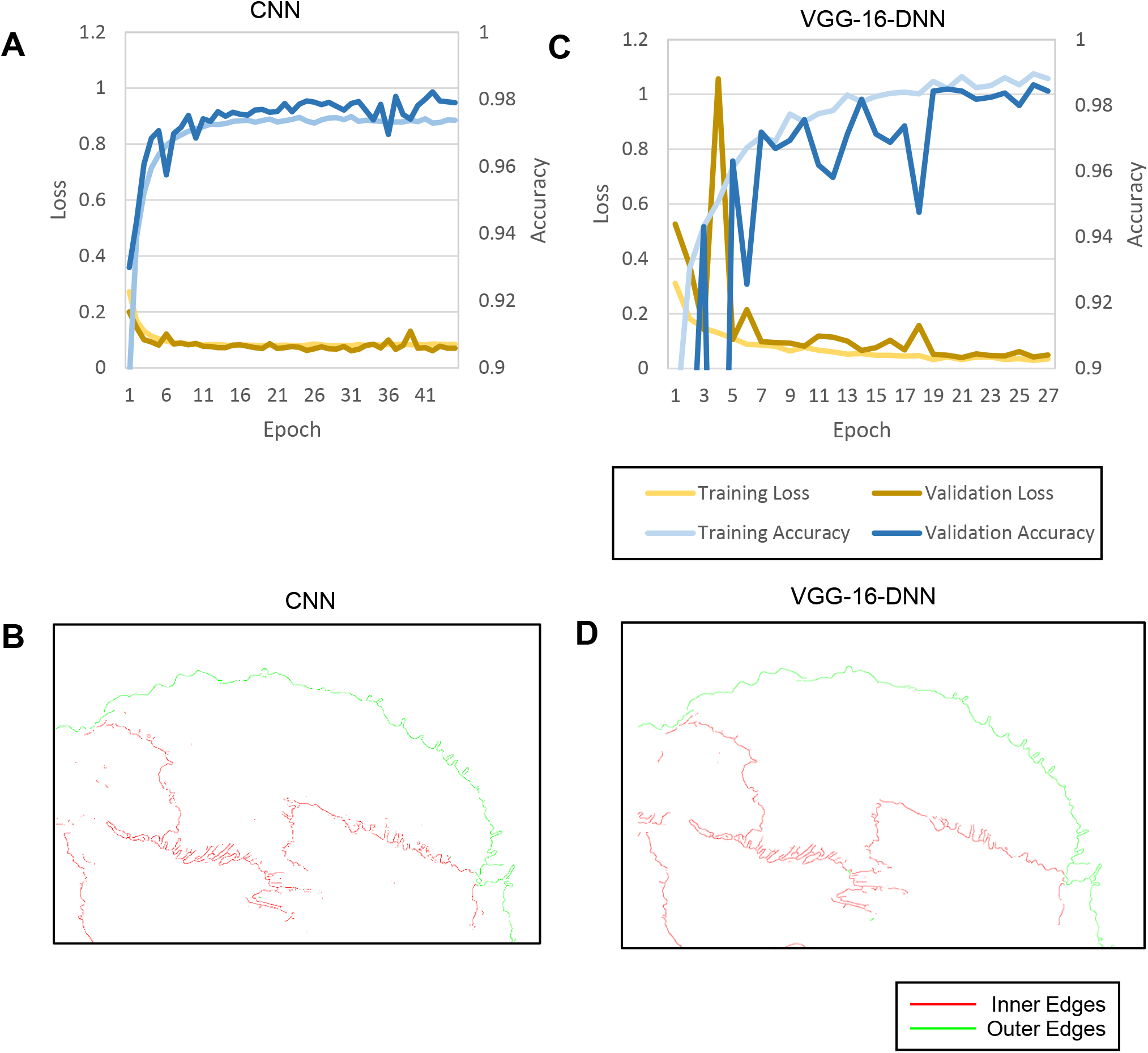
Deep Learning Training Results. (**A**) The training curve represents a training progress with CNN. (**B**) Edge image reconstruction based on the results of (B). (**C**) The training curve represents a training progress with VGG-16-DNN. (**D**) Edge image reconstruction based on the results of (C).

In order to reduce the amount of the training set, the data augmentation and dropout were included in the CNN training. To see their effects on the training performance, we compared the results using Dice Similarity Coefficients indicating the similarity between the edges from deep learning and the ground truth^36^. We chose the testing frame #16 which is in the middle of the training frame #1 and #31, and the testing frame #46 in the middle of the training frame #31 and #61. We also chose the testing frame #191 in order to see how trained edge detection efficiency decreased when the testing image is not close to the training images. As in Fig. 5A, the data augmentation and dropout were highly effective in substantially increasing the Dice Similarity Coefficients and helped the CNN to find some of the missing edges in the images (Fig. 5F). The VGG-16 pretrained model also further increased the Dice Similarity Coefficients in all testing frames and allowed the DNN to produce visually better results (Fig. 5G).

**Figure 5:**
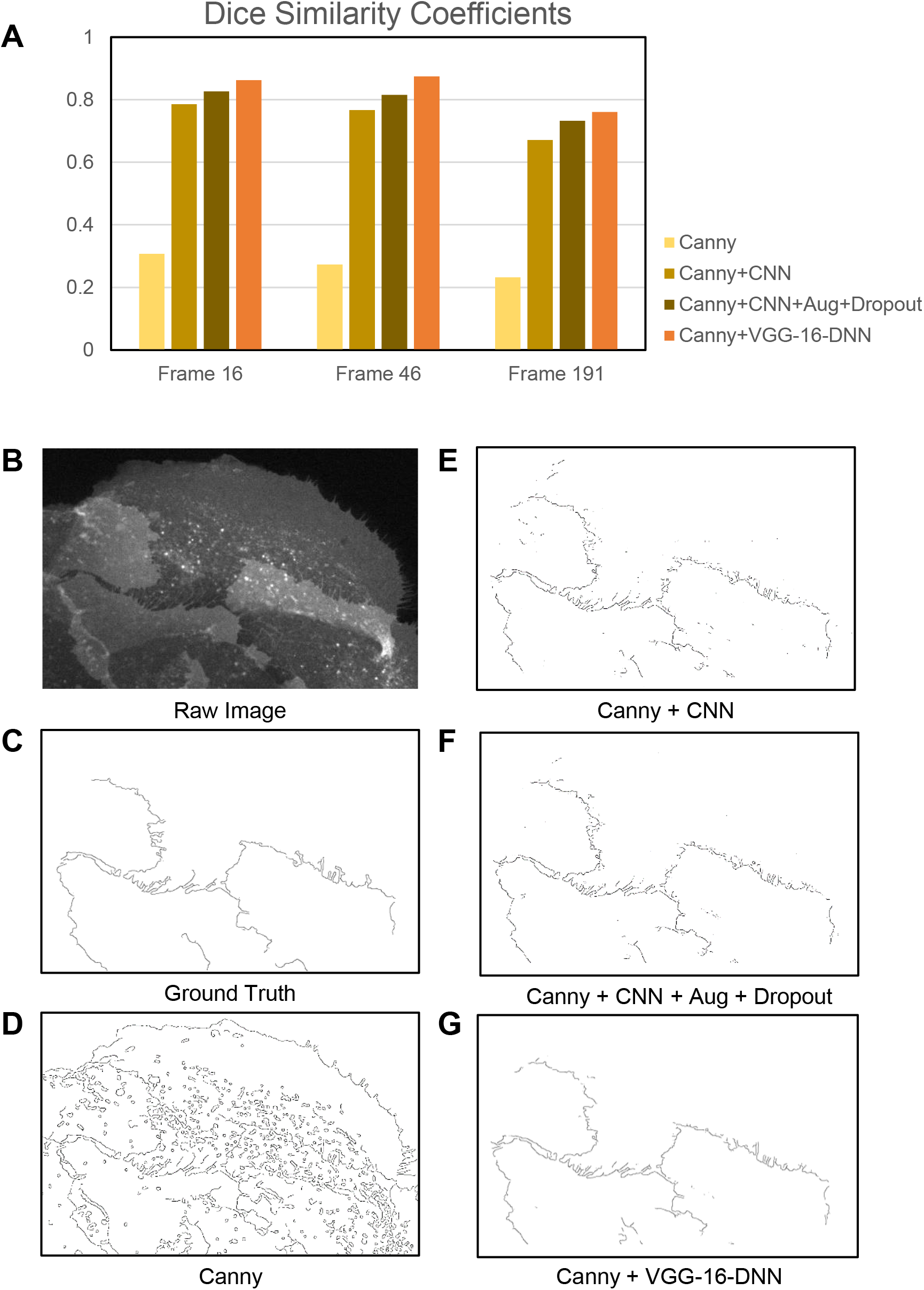
Roles of augmentation and dropout for testing performance. (**A**) Dice similarity coefficients in different combination of algorithms in each test frame (#16, 46, and 191) (**B**) Raw image. (**C**) Ground truth of the inner edges. (**D**) Canny image. (**E**) Canny + CNN. ( **F**) Canny + CNN + Augmentation + Dropout. ( **G**) Canny + VGG-16-DNN.

After we had final images of edges from the VGG-16-DNN, we ran post-processing to remove misclassified edges. We measured the pixel numbers of each edge. If this number was less than 5 pixels, we treated them as noises and discarded them. This process was able to clean up some noises within the final image (Fig. 6A). Then the local thresholding was applied to the image patches around the detected edges based on the median intensity of the edges within the patches (Fig. 6B). This operation tends to produce a large segmented object with real edges. Therefore, the large segmented objects were selected and the boundary was refined by the image close operation to fill the small gaps within the masks, followed by edge extraction (Fig. 6C-D). As demonstrated, the final results were highly accurate edge detection of inner edges of cryptic lamellipodia (Fig. 6E-F), and the time evolution of the cryptic lamellipodial edge was obtained as in Fig. 6G.

**Figure 6:**
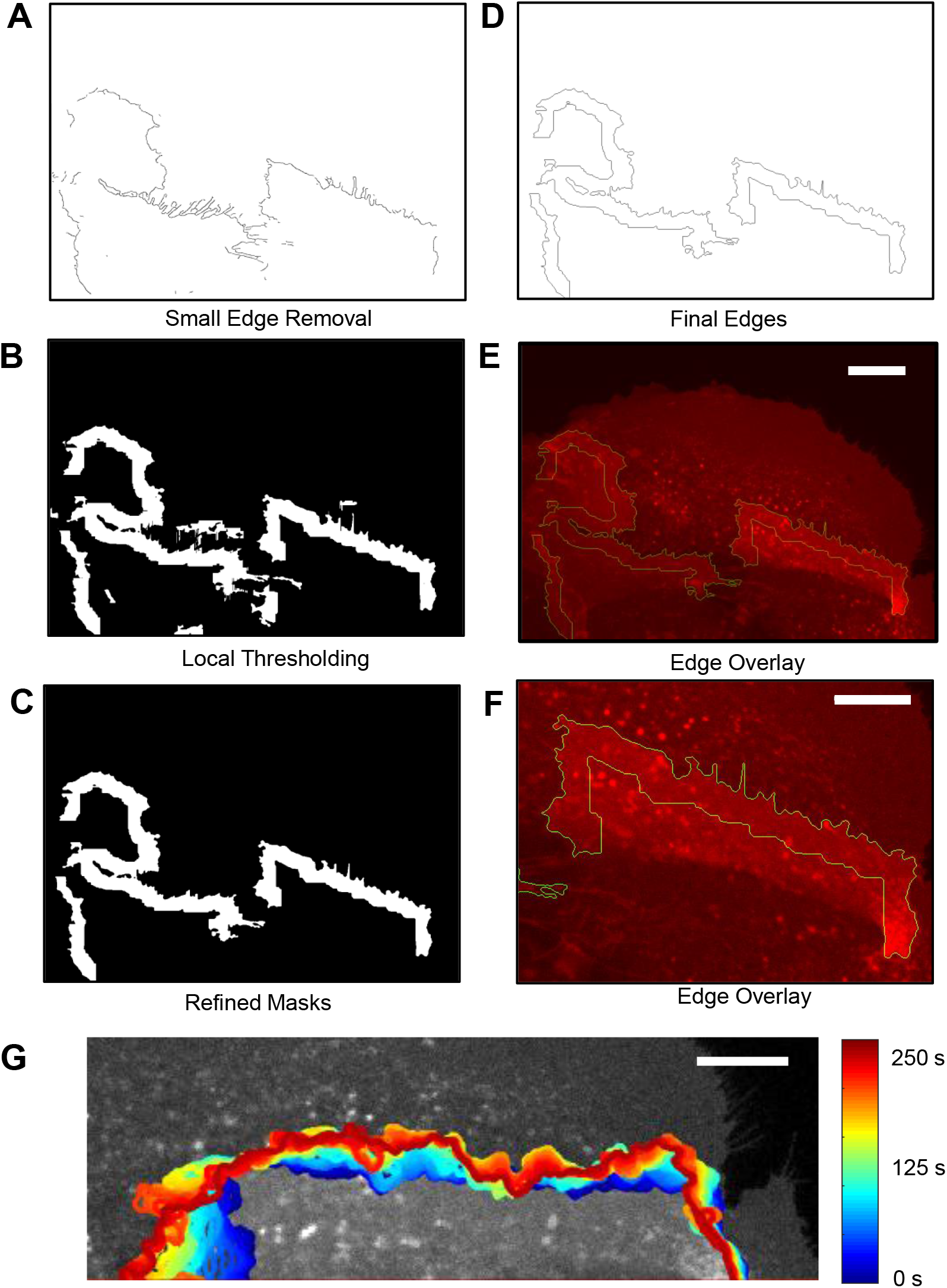
Results of Local Thresholding Based on the Edge Classification by VGG-16- DNN. (**A**) The small edges (< 5 pixels) are discarded. (**B**) The locally thresholded image. (**C**) The refined segmented images where the small segmented areas were discarded and the edge was cleaned by mask open/close operation. (**D**) the edge images of (C). (**E**) Final edge image (Green) overlayed with the raw intensity image (Red). Bar: 15mm. (**F**). The magnified images of (E). Bar: 10mm. (**G**) Edge evolution of (F). The image was rotated for better visualization. Bar: 10mm.

## Conclusion

This work serves to create an efficient method of detecting cell edges from cryptic lamellipodia in cell images for collective cell migration studies. By assisting the conventional image analysis algorithms using deep learning techniques, our pipeline was shown to detect cell edges at a high performance. Whereas most of the deep learning application to image segmentation is completely independent of the conventional algorithms^24,25^, we aimed to take advantage of deep learning to enhance the conventional approaches. The advantage of this deep learning application is that it allows us to take advantage of decades of works in computational image analysis without replacing them. In this work, we demonstrated that deep learning is flexible enough to be incorporated into existing analysis workflow and substantially improve the analysis results.

Using Canny edge detection on the original cell image, we confined training within the edge candidate points suggested by the Canny edge detector. This simplified the training, reduced the computation time, and required less data for the deep learning. Furthermore, the inner edges suggested by the CNN provided important information with the local thresholding algorithms to achieve highly accurate continuous edges. This work demonstrated the feasibility of deep learning application in complex cell segmentation or cell migration studies.

One of drawbacks of deep learning is that it generally requires a large training set to prevent overfitting. In order to address this issue, we also tested various techniques known to mitigate overfitting with limited data. We applied data augmentation, dropout, and the pretrained model, VGG-16. Using these techniques, we were able to train DNNs using the training data every 25 frames, which makes it practical to apply deep learning techniques to live cell image segmentation.

Images analyses highly rely on the accuracy of edge detection. However, when cells overlap with each other, it is increasingly difficult to robustly detect cell edges due to the complex background patterns. Usually conventional algorithms alone are unable to reliably distinguish cell edges in an image where cells are overlapping. Therefore, the computational studies of cell biological phenomena during intercellular interaction are limited. This pipeline showed a beneficial application that can be made to the research on cryptic lamellipodial protrusion in collective cell migration. We believe that this pipeline is not limited in the cryptic lamellipodial segmentation. It can be also used in many time-lapse movies where cells interact with neighboring cells.

## Acknowledgements

We thank NVIDIA for providing us with TITAN X GPU cards (NVIDIA Hardware Grant Program), Microsoft for providing us with Azure cloud computing resources (Microsoft Azure Research Award), and Boston Scientific for providing us with the gift for deep learning research. This work is supported by the WPI Start-up Fund for new faculty.

## Author Contributions

C. W. initiated the project, designed the algorithm of local thresholding, and wrote the final version of the manuscript and supplement. S. K and E. K. performed training of DNNs and wrote the final version of the manuscript and supplement. X. Z applied the pretrained model. H. C. performed the fluorescence live cell imaging and wrote the final version of the manuscript and supplement; A. C. contributed to the writing of the manuscript; K. L. coordinated the study and wrote the final version of the manuscript and supplement. All authors discussed the results of the study.

## Author Information

The authors declare no competing financial interests. Correspondence and requests for materials should be addressed to K.L.

